# EFA6R suppresses ovarian cancer cell migration and invasion

**DOI:** 10.1101/2022.01.21.477266

**Authors:** Salman Tamaddon-Jahromi, Kate Murphy, William Walker, Venkateswarlu Kanamarlapudi

## Abstract

Exchange factor for ADP-ribosylation factor (Arf)6 (EFA6)R expression loss in ovarian cancer has shown to decrease patient survival. EFA6R contains the catalytic Sec7, pleckstrin homology (PH), and coiled-coil (CC) domains. To gain further insight into the role of EFA6R, this study further investigated EFA6R expression in OC and its putative role as a metastatic suppressor. EFA6R mRNA expression, assessed by RT-qPCR, was significantly downregulated in OC tissues and cell lines. OC tissue microarray staining with EFA6R antibody showed that loss of protein expression correlated with increased cancer grade. Furthermore, EFA6R protein levels, assessed by immunoblotting, were significantly reduced in OC tissues and cell lines. Treatment of SKOV-3 cells with 5-aza-2’deoxycytidine, an epigenetic regulator, restored EFA6R expression and attenuated functional cell migration and invasion, which was reversed by siRNA-mediated knockdown of EFA6R expression. This study also revelated that exogenously expressed EFA6R localises to the plasma membrane, through its PH domain, and thereby inhibits cell migration and invasion in the CC domain-dependent and an Arf6-independent manner. EFA6R loss-of-function involves epigenetic mechanisms in which downregulation increases OC tumour cell migration and invasion.

**Summary statement:** EFA6R expression is epigenetically regulated in ovarian cancer cells and loss of expression correlates with increased tumour grade and enhanced tumour cell migration. EFA6R appears to mediate these effects independently of Arf6.

## INTRODUCTION

Ovarian cancer is the most lethal gynaecological cancer in women (1). Epithelial OC is the most common form of OC. Cancer metastasis and the acquired resistance to drug treatment are the major causes of ovarian cancer-associated deaths (1). OC is classified into stages 1-4 based on no metastasis (stage 1), metastasis of the primary tumour to the pelvic cavity (stage 2), abdomen wall (stage 3) or distal metastasis (stage 4). OC subtypes include serous (from the fallopian tube), endometrium (from the endometroid), mucinous (from the endocervix) and clear cell (within the vagina). Our understanding of the impact of metastatic suppressors in OC is currently limited, therefore identification of novel metastatic regulators can potentially serve as prognostic biomarkers, therapeutic targets and predictors of treatment response.

Exchange factor for ADP-ribosylation factor (Arf)6 (EFA6)R expression loss in OC has been linked to a decrease in patient survival (2). EFA6R is a member of the EFA6 family of guanine exchange factors (GEFs), which activate Arf6 small GTPase (3). Arf6 small GTPase mediates membrane trafficking and cytoskeleton reorganisation at the plasma membrane by cycling between the active GTP-bound and inactive GDP-bound forms (4). Arf6 and its regulators not only show altered expression in many cancers but also promote cancer metastasis and drug resistance (5). EFA6R shares a common domain organisation with the other members of the EFA6 family and the wider Arf6 GEFs (3,6). It has a Sec7 catalytic domain which preferentially activates Arf6, a phosphatidylinositol 4,5-bisphosphate (PI4,5-P2) binding pleckstrin homology (PH) domain and a coiled-coil (7) region, which is both necessary for an efficient plasma membrane localisation and F-actin re-organisation (8). In addition to its role in OC, EFA6R expression is altered in several other cancers (3,8). Another member of the EFA6 family, EFA6B, has also been shown to antagonise breast cancer by promoting tight junction (TJ) proteins claudin-2 and occludin expression as well as blocking the transformer growth factor-beta pathway through Arf6 activation (7).

In OC, a functional role for EFA6R has yet to be identified, although downregulation of EFA6R expression has previously been shown to have a drastic impact on patient survival (2). Aberrant gene loss in OC is interconnected with epigenetic alterations, where DNA hypermethylation and histone de-acetylation within or upstream of promoter regions of tumour suppressor genes (TSGs) have been shown to lead to undesirable gene silencing (9). Therefore, epigenetic inhibitors have been utilised to revive the expression of many TSGs, subsequently reversing adverse phenotypes (10). Here, we evaluated EFA6R expression in OC and established a correlation between EFA6R downregulation and OC metastasis.

## RESULTS AND DISCUSSION

### EFA6R expression is downregulated in OC

It has been shown previously that EFA6R is amongst a cluster of genes on chromosomal region 8p22, whose expression loss in OC has a detrimental impact on patient survival (2). To confirm this, we initially assessed *EFA6R* mRNA expression in the OC using an ovarian TissueScan™ cDNA array. When compared with that in healthy ovarian tissue, *EFA6R* mRNA expression was significantly reduced in cancers of grade I, and more markedly reduced in cancers of grade II and III (Fig. 1A). Given the heterogeneity in histologic origin, sensitivity to treatment and differential prevalence of OC subtypes, we also analysed the *EFA6R* mRNA expression based on the OC subtype (Fig. 1B). This revealed a significant reduction of *EFA6R* levels in all (endometroid, clear cell and serous) but the mucinous subtype of OC. This is most likely due to a limited sample cohort, but we do not rule out a biological reason for this observation. Previous studies have shown that mucinous carcinomas differ from the other subtypes of OC in the frequency and pattern of loss of heterozygosity (LOH) at 8p (11).

**Figure 1.**
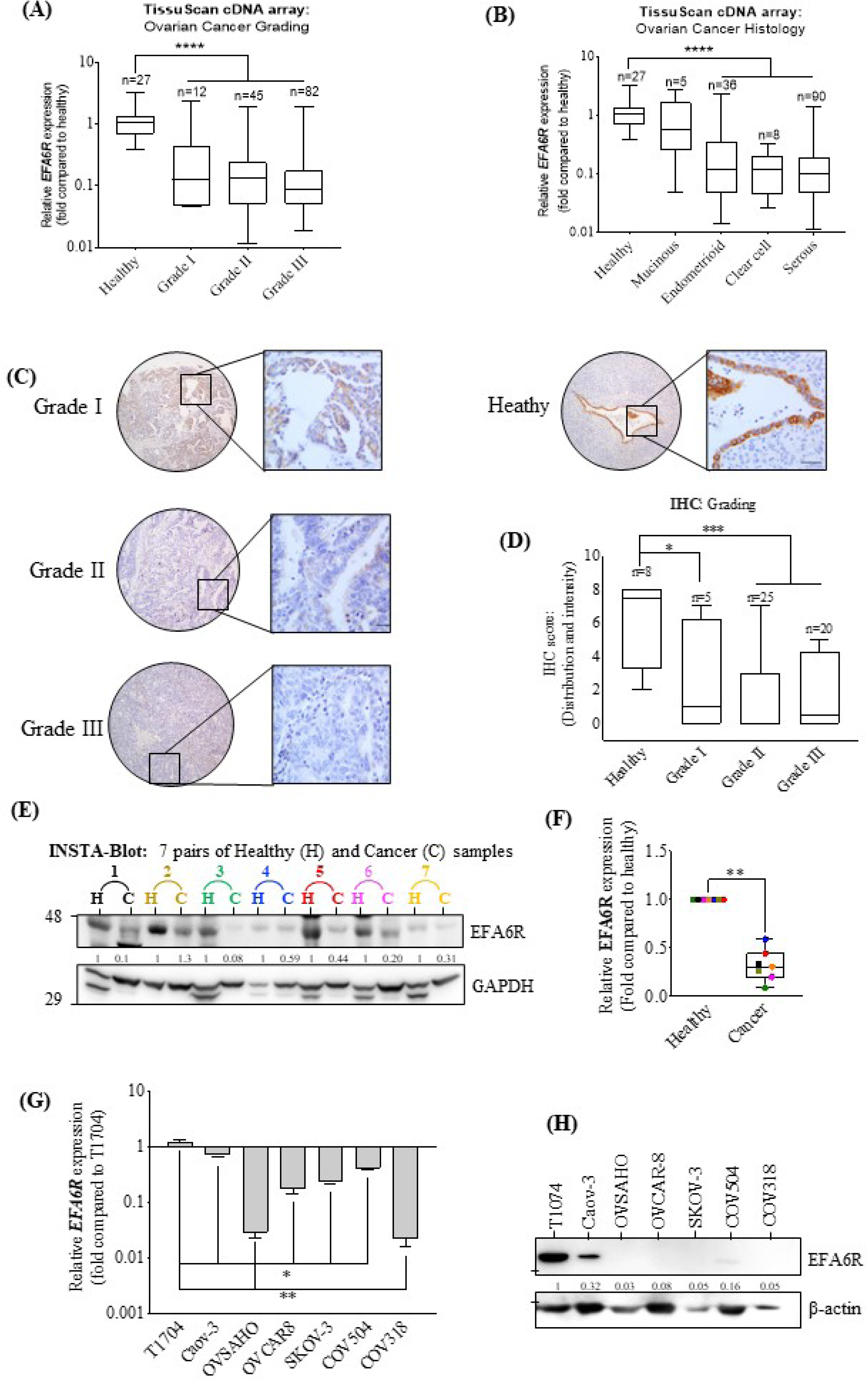
EFA6R expression is significantly reduced in OC. **A** RT-qPCR analysis of *EFA6R* mRNA expression in human ovarian healthy and cancer tissues and the results displayed based on OC histology grading. **B** The data in (**A**) was separated based on four OC subtypes: mucinous, endometroid, clear cell and serous. The box plots in (**A**) and (**B**) show the median fold-change (\*\**P* <0.01, \*\*\**P* <0.001, and \*\*\*\**P* <0.0001, based on Kruskal–Wallis test with Dunn’s test). **C** Representative immunohistochemical (IHC) images of the EFA6R protein expression in a TMA containing human ovarian healthy and OC tissue sections, probed with an anti-EFA6R antibody. The scale bar is 100μm. **D** The IHC score, based on total sum of distribution of stain (from 0 to +5) and stain intensity (0 = none, +1 = weak, +2 = moderate and +3 = strong). The box plots show the median IHC score (\**P* <0.05, \*\*\*\**P* <0.0001, based on Kruskal–Wallis test with Dunn’s test). **E** Western blot analysis of EFA6R expression in OC tissue and healthy adjacent ovarian tissue. An ovarian cancer OncoPair INSTA-Blot probed with the anti-EFA6R and anti-GAPDH (loading control) antibodies. **F** Pooled fold-change in EFA6R protein expression, as assessed by Western blot in (**E**), in healthy tissue versus OC tissue (\*\**P* <0.01, based on one-sample t-test). **G** RT-qPCR analysis of *EFA6R* mRNA expression in human ovarian non-malignant (T1704) and OC cell lines. The data are presented as the mean ± s.e.m. from three independent experiments (n = 3) (\**P* <0.05, \*\**P* <0.01, \*\*\*\**P* <0.0001, based on Mann-Whitney test). **H** Western blot analysis of EFA6R protein expression in ovarian non-malignant (T1704) and OC cell lines using the anti-EFA6R and anti-ß-actin (loading control) antibodies. The intensity of the bands was quantified by using Image J software, normalised to the expression of ß-actin and shown below the EFA6R blot.

To further evaluate the relative expression of EFA6R at the protein level, immunohistochemical staining was performed on an ovarian tissue microarray (TMA) (Fig. 1C). Based on the average total sum of EFA6R immunostain distribution and intensity, EFA6R protein expression in the TMA was quantified (Fig. 1D). Overall, there was a significant loss in EFA6R expression when compared between healthy tissue and Grades I-III cancer cases. In further analysis, the expression of EFA6R protein in the lysates of tissues obtained from OC patients and their healthy counterparts were assessed by immunoblotting (Fig. 1E). EFA6R protein expression was significantly reduced in tumour sample lysates when compared to that from healthy tissue. Using available OC cell lines, EFA6R mRNA expression in malignant cell lines was compared with that in the ovarian non-tumour cell line (T1074) (Fig. 1G, 1H). Compared to that in non-malignant cells, *EFA6R* mRNA expression was significantly reduced in OC cell lines. Furthermore, when assessed by immunoblotting, EFA6R protein expression was either undetectable or markedly reduced in the majority of OC cell lines. Overall, EFA6R expression loss is evident during the early stages of OC development, suggesting that EFA6R could be a useful biomarker for OC tumours.

### Restoration of EFA6R expression attenuates OC cell migration and invasion

Inactivation of gene expression through hypermethylation, by DNA methyltransferases (DNMTs), and histone de-acetylation, by histone de-acetylases (HDACs), has previously been described for several TSGs in OC (12-14). Therefore, using an OC cell line (SKOV-3), we tested whether suppression of EFA6R expression is associated with either of these epigenetic mechanisms. We began by assessing SKOV-3 sensitivity to the DNMT inhibitor 5-Aza-CdR, or the HDAC inhibitor suberanilohydroxamic acid (SAHA) by treating cells with 0.1-10μM of 5-Aza-CdR or SAHA for four days and assessing the viability (Fig. 2A). Although 5-Aza-CdR was relatively non-toxic, a significant reduction in SKOV3 cell viability was observed with 10μM SAHA. Treatment of SKOV-3 cells with 10μM 5-Aza-CdR, but not 1.0μM SAHA, significantly restored EFA6R protein expression (Fig. 2B), suggesting that DNA hypermethylation plays a major role in the epigenetic silencing of EFA6R gene expression. Therefore, in subsequent functional assays, we used 10μM 5-Aza-Cdr treatment to re-establish EFA6R expression.

**Figure 2.**
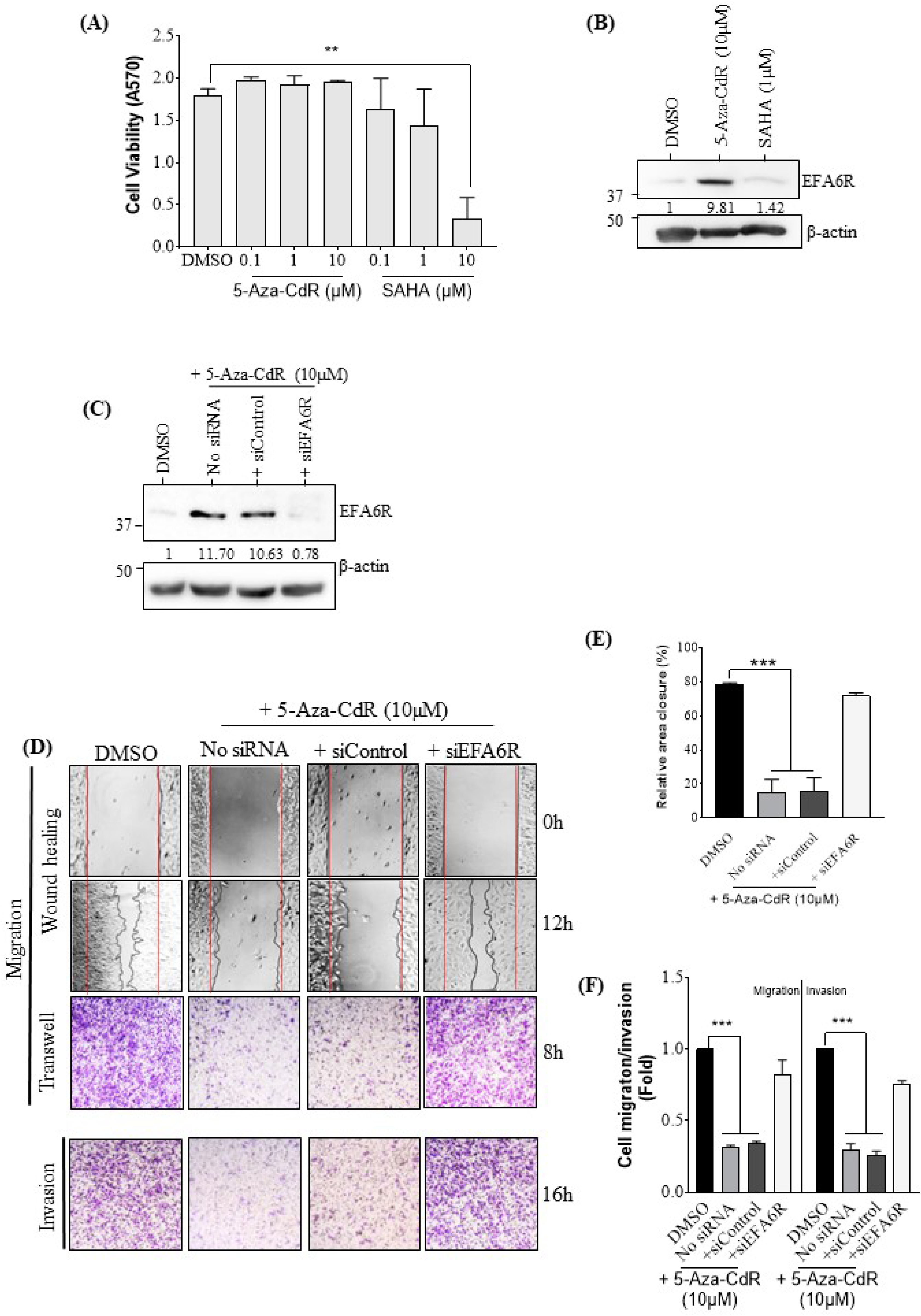
EFA6R suppresses SKOV-3 cell migration and invasion. **A** Dose-dependent effect of epigenetic drugs (5-Aza-CdR and SAHA) on SKOV-3 cell viability (displayed as absorbance at 570nm [A570]). The data are shown as mean ± s.e.m., n=3, \*\**P* <0.01, One-way ANOVA with Dunnett’s test. **B** Western blot analysis of EFA6R protein expression in SKOV-3 cells following a 4-day treatment with 10μM 5-Aza-CdR or 1.0μM SAHA (DMSO = solvent control); probed with the anti-EFA6R and anti-β-actin (loading control) antibodies. **C** Western blot analysis of EFA6R protein expression in SKOV-3 cells, following a four-day treatment with 10 μM 5-Aza-Cdr with no siRNA or transfected with siControl or siEFA6R (DMSO = solvent control); probed with anti-EFA6R and anti-β-actin (loading control) antibodies. The intensity of the bands was quantified, normalised to the expression of ß-actin and presented below the EFA6R blot as fold change. **D** Representative images of SKOV-3 cell migration (assessed by using wound healing and transwell migration assays) and invasion (assessed by using transwell coated with Matrigel). In the wound healing assay, cell migration is shown at 0h and 12h following removal of the Ibidi insert. Cell migration was also assessed using a transwell migration assay. The bottom panel shows cell invasion assessed using Matrigel-coated transwell migration inserts. Following 8h of cell migration and 16h of cell invasion, the cells were fixed with 4% PFA and stained with 0.2% crystal violet. **E** Relative cell migration was assessed in wound healing assay by measuring gap closure (%) compared to that in control post-12h of the Ibidi insert removal. The data are presented as mean ± s.e.m. (n=3, \*\*\**P* <0.001, based on One-way ANOVA with Dunnett’s test). **F** The crystal violet staining was extracted from migrated or invaded cells using 5% SDS and its absorbance was measured at 570nm. The data are shown as mean ± s.e.m. (n=3, \*\*\**P* <0.001, based on One-way ANOVA with Dunnett’s test).

5-Aza-Cdr restored EFA6R protein expression could be knocked down with an EFA6R-specific siRNA (siEFA6R) (Fig. 2C). Since cell migration and cell invasion are key steps in cancer metastasis, we next assessed the effect of restoration of EFA6R expression on cell migration and invasion by using wound healing and transwell migration/invasion assays (Fig. 2D). In these assays, SKOV-3 cells treated with 5-Aza-Cdr plus control siRNA exhibited reduced migration and invasion. In comparison, solvent (DMSO)-treated cells or cells concomitantly treated with 5-Aza-Cdr and siEFA6R exhibited increased cell migration (Fig. 2E). Comparative analysis showed that 5-Aza-Cdr treatment produced a ∼3-fold reduction in both cell migration and invasion and this was effectively reversed by siEFA6R treatment (Fig. 2F). Overall we demonstrated that restoration of EFA6R expression leads to a significant reduction in both cell migration and invasion. This reduced metastatic phenotype was specifically related to the re-establishment of EFA6R expression in ovarian cancer cells, as siRNA-mediated knockdown of EFA6R reversed this effect. Loss of EFA6R expression may, therefore, play a significant role in OC metastasis.

### EFA6R suppresses cell migration/invasion in an Arf6 GEF activity-independent manner

To determine whether plasma membrane localisation or the GEF activity or both are required for EFA6R to suppress cell migration and invasion, we analysed the effect of GFP-tagged EFA6R and its deletion mutants overexpression on SKOV-3 cells cell migration and invasion (Fig. 3A). EFA6R and the Sec7 deletion mutant (GFP-EFA6R ΔSec7) proteins showed expected plasma membrane localisation due to the presence of intact PH and CC domains (Fig. 3B). However, the deletion mutants of EFA6R lacking either the PH domain (GFP-EFA6R ΔPH) or the CC domain (GFP-EFA6R ΔCC) showed absent or weak (respectively) plasma membrane localisation. The addition of the CAAX motif of K-Ras at the C-terminus of EFA6R without the PH (GFP-EFA6R ΔPH_CAAX_) or CC domain (GFP-EFA6R ΔCC_CAAX_) restored the plasma membrane localisation of EFA6R (8). Immunoblotting confirmed the similar expression of GFP-tagged EFA6R and its deletion mutants in SKOV-3 cells post-transfection (Fig. 3C).

**Figure 3.**
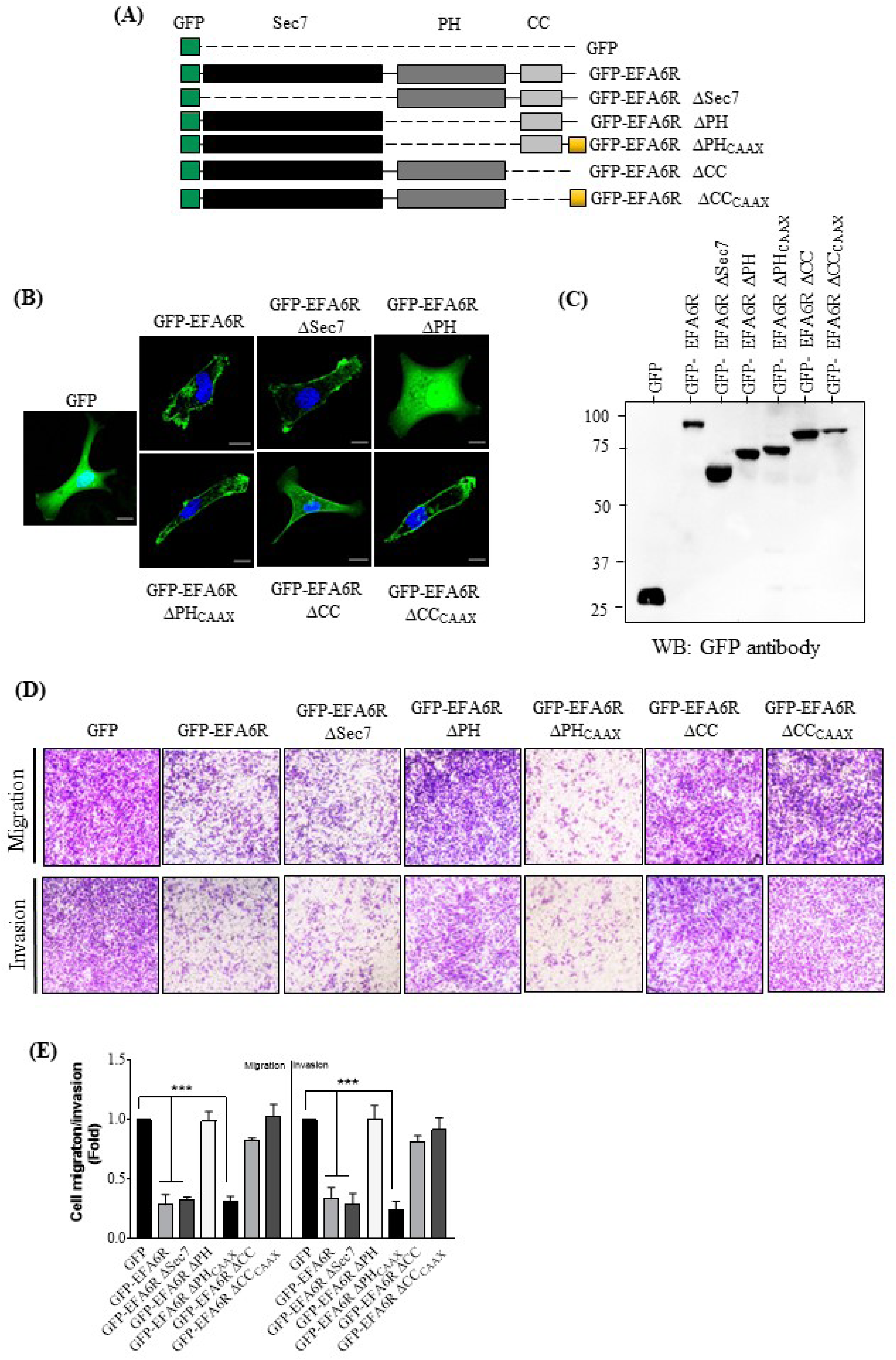
EFA6R attenuates cell invasion in a CC domain-dependent manner. **A** Schematic view of GFP and GFP-tagged EFA6R and its deletion constructs used in this study. **B** Analysis of intracellular localisation of GFP-tagged EFA6R and its deletion constructs expressed in SKOV-3 cells using confocal fluorescence microscopy. Images are representative of 75-100 cells. **C** Western blot (WB) analysis of GFP-tagged EFA6R and its deletion constructs expression in SKOV-3 cells by using an anti-GFP antibody. **D** Representative images of migration and invasion (assessed using Transwell inserts) of SKOV-3 cells transfected with GFP-tagged EFA6R and its deletion mutants. 8h post-migration and 16h post-invasion, the cells were fixed with 4% PFA and stained with 0.2% crystal violet. **E** Crystal violet staining of migrating/invading cells was extracted using 5% SDS and the absorption was measured at 570nm. The data are presented as mean ± s.e.m. (n=3, \*\*\**P* <0.001, based on One-way ANOVA with Dunnett’s test).

We further determined the effect of EFA6R and its deletion mutants on SKOV-3 cell invasion and migration (Fig. 3D). As expected, the migratory and invasive abilities of SKOV-3 cells expressing GFP-EFA6R were significantly reduced when compared to that of control GFP-transfected cells (Fig. 3E), confirming the anti-metastatic role of EFA6R in SKOV-3 cells. Expression of the GFP-EFA6R ΔSec7 also reduced cell migration/invasion, similar to that seen with GFP-EFA6R. Therefore, the Sec7 domain of EFA6R, which is the GEF domain required for the activation of Arf6, is not essential for attenuation of cell migration or invasion, indicating that EFA6R regulates OC cell metastasis independently of the Arf6 pathway. Indeed, consistent with this finding, EFA6R-mediated inhibition of cell migration/invasion was not reversed by siRNA-mediated down-regulation of Arf6 expression (siARF6) (Fig. 4). In contrast, EFA6R ΔPH, which lacks plasma membrane localisation, failed to inhibit SKOV-3 cell migration/ invasion and this was reversed by targeting it to the plasma membrane by adding the CAAX motif (EFA6R ΔPH_CAAX_), suggesting that the PH domain of EAF6R does not play a functional role in the attenuation of cell migration and invasion. Interestingly, we observed that EFA6R-mediated inhibition of cell migration/invasion was reversed by either removal of the CC domain (GFP-EFA6R ΔCC) or removing the CC domain and adding the CAAX domain (GFP-EFA6R ΔCC_CAAX_). This indicates that the CC domain plays an important role in the EFA6R-mediated attenuation of cell migration and invasion in SKOV-3 OC cells.

**Figure 4.**
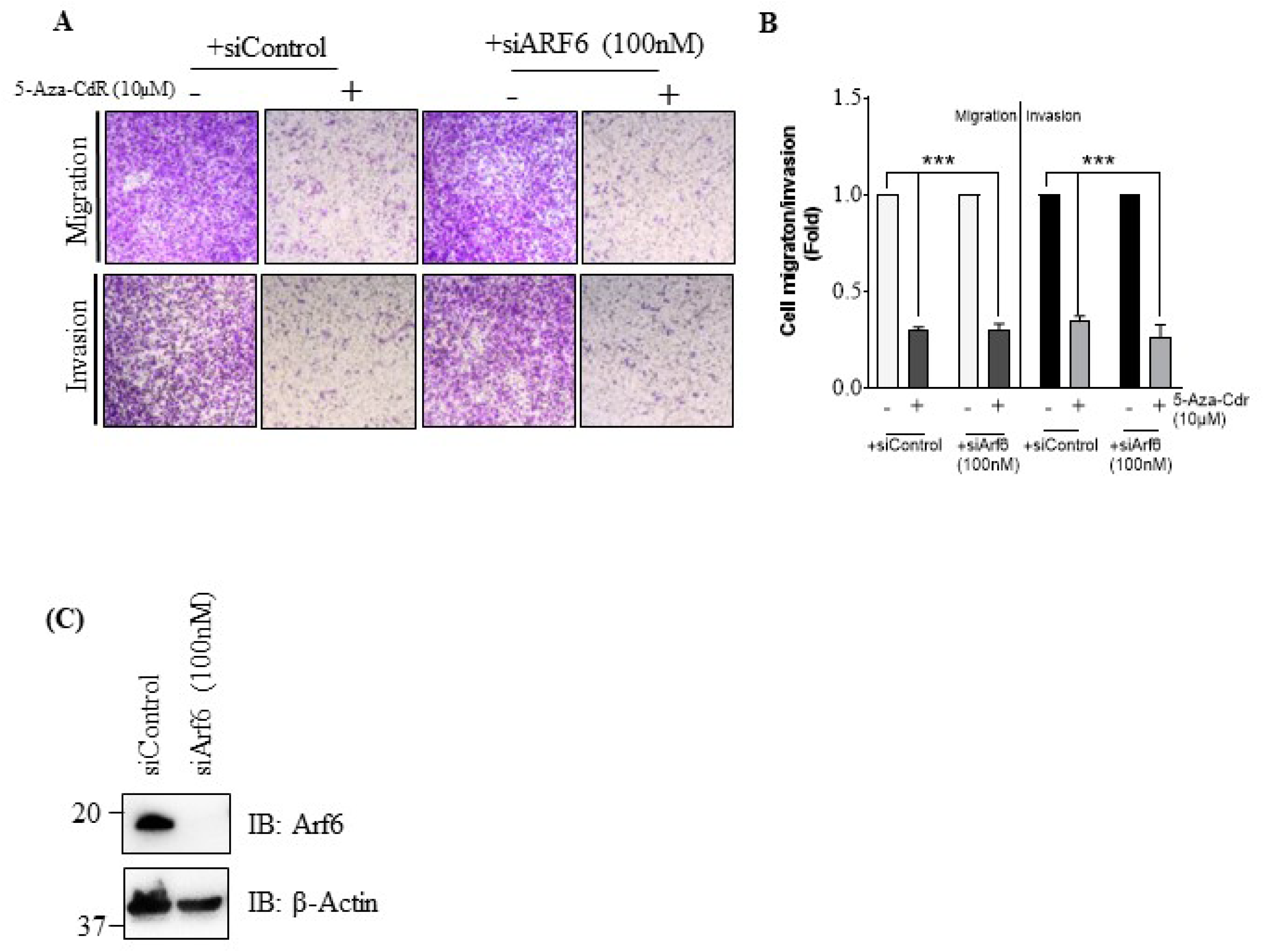
SKOV-3 cell migration and invasion are independent of the Arf6 pathway. **A** Representative images of SKOV-3 cell migration and invasion (assessed by using transwell inserts). Following a 4-day treatment with 10μM 5-Aza-Cdr with no siRNA or transfected with siControl or siArf6, SKOV-3 cells were subjected to cell migration and invasion (using transwell insert). 8h post-migration and 16h post-invasion, the cells were fixed with 4% PFA and stained using 0.2% crystal violet. **B** The crystal violet staining of invading cells was extracted using 5% SDS and the absorption was measured at 570nm. Errors bars represent the ± s.e.m. (n=3, \*\*\**P* <0.001, based on One-way ANOVA with Dunnett’s test). **C** Western blotting analysis of Arf6 expression in SKOV-3 cells transfected for 4 days with 100nM siControl or siArf6 using an anti-Arf6, an anti-ARF1 and an anti-β-actin (loading control) antibodies.

We show that cell migration/invasion occurs independently of the Arf6 pathway, as deletion of the Sec7 domain or siRNA-mediated downregulation of Arf6 did not affect these cellular activities. These results were particularly surprising since the role of Arf6 in cancer cell migration and invasion has been extensively documented in other cancers, including breast, lung and pancreatic carcinomas (5,15-21). EFA6R localisation to the plasma membrane, through the PH domain, is important for its ability to inhibit cell migration and invasion. However, the PH domain itself is functionally irrelevant to the inhibition of cell invasion by EFA6R, as its requirement for the membrane localisation of EFA6R can be bypassed using the CAAX tag. Furthermore, it is notable that the CC domain plays an important functional role in attenuating cell migration and invasion and the CAAX motif did not bypass the functional importance of the CC domain. Taken together, these findings demonstrate that the CC domain of EFA6R may be the site of possible protein interactions, by which it negatively regulates cell metastasis. In contrast, the PH and Sec7 domains do not seem to play a functional role in the inhibition of cell migration or invasion. Indeed, the CC domain of EFA6R has previously been shown to be responsible for cytoskeleton rearrangements and interactions with downstream signalling proteins (8), however, no interacting proteins of the EFA6R CC domain have been identified so far.

In summary, we demonstrated here that EFA6R is epigenetically suppressed in OC. DNA hypermethylation is particularly involved in that the DNMT inhibitor (5-Aza-CdR) restored EFA6R expression in OC cells, thereby reducing their ability to migrate and invade. Furthermore, EFA6R-mediated cell migration and invasion occurs in an Arf6-independent manner and appears to be mainly regulated through its CC domain, highlighting the need for further investigations into the exact function of this EFA6R domain in OC cells. Further studies into the suitability of using EFA6R expression as an early detector of OC may prove to be transformative for OC diagnosis. OC can be cured in more than 90% of cases if it is detected early.

## MATERIALS AND METHODS

### Plasmids

The GFP-EFA6R, GFP-EFA6R ΔSec7, GFP-EFA6R ΔPH and GFP-EFA6R ΔCC expressing plasmids used in this study have been described previously (8). The GFP-EFA6R ΔPH and GFP-EFA6R ΔCC were targeted to the membrane (GFP-EFA6R ΔCC_CAAX_ and GFP-EFA6R ΔPH_CAAX_) by attaching a C-terminal *CAAX* motif using polymerase chain reaction (PCR) with a 3’-primer containing the coding sequence for the CAAX motif from K-Ras as described previously (22). Human EFA6R siRNA (5′-GCUACUGAGUAACGAUGAA-3′) and negative control siRNA (siControl) have also been described previously (8).

### Cell culture and transfection

Human ovarian non-malignant (T1074) and malignant (SKOV-3) cell lines were provided by Prof. Deyarina Gonzalez (Swansea University). OVSAHO and Caov-3 cell lines were provided by Dr Marion Curtis (University of Chicago). OVCAR8, COV504 and COV318 cell lines were provided by Dr Alan Richardson (Keele University). All cell lines were cultured aseptically at 37°C/5% CO_2_ in RPMI 1640 (Sigma, UK) supplemented with 10% foetal bovine serum (FBS; Sigma, UK), 100 U/ml penicillin, 0.1 mg/ml streptomycin (Sigma, UK) and 2 mM GlutaMAX (Gibco, UK). Cell counting was performed using a Countess cell counting chamber slide (Invitrogen, UK). SKOV-3 cells were transiently transfected with either 100nM siRNA or 2μg plasmid DNA by electroporation using the Neon transfection system (Thermo Fisher Scientific, UK) using the following parameters: 1170 pulse voltage, 30ms pulse width and 2 pulse number. Following four days of incubation, the transfected cells were subjected to various functional assays. The transfection efficiency of the EFA6R and its deletion constructs was assessed by measuring GFP-derived fluorescence (of GFP-tagged EFA6R and its deletion mutants) using standard flow cytometry (23).

### Real-Time Quantitative Polymerase Chain Reaction (RT-qPCR)

*EFA6R* gene (mRNA) expression was carried out in healthy and cancer ovarian tissues by RT-qPCR using Tissuescan™ ovarian cancer cDNA arrays I-IV (Origene Technologies, USA) and EFA6R (NM_206909.3) sequence-specific primers (Forward 5’-CGCAGCGGCAGAGACATTT-3’ and Reverse 5’-TTTGGCCTTGGCAACACTCT-3’); the cDNA arrays were pre-normalised to β-actin expression. For cell lines, total RNA was extracted from cell lines using TRI Reagent (Sigma, UK). cDNA was prepared from total RNA using the High Capacity cDNA Reverse Transcription Kit (Thermo Fisher Scientific, UK) (24). RT-qPCR was carried out using the EFA6R primers (described above), β-actin as housekeeping gene (Forward: 5’-CAGCCATGTACGTTGCTATCCAGG-3; reverse: 5’-AGGTCCAGACGCAGGATGGCA-3’) and 2X RT^2^ SYBR Green qPCR master mix (Qiagen, UK). The relative fold change in gene expression was analysed using the double delta Cq analysis (2^-ΔΔCq^) method (25).

### Western blotting

Total protein was extracted from cell lines using TRI Reagent (Sigma, UK). 40μg of isolated protein samples were then separated using 10% sodium dodecyl sulphate-polyacrylamide gel electrophoresis (SDS-PAGE) for 40min at 200 volts, then transferred, using Trans-blot Turbo transfer system (Bio-Rad, UK), on to polyvinylidene fluoride (PVDF) membrane (GE Healthcare Life Sciences, UK) at 25 volts for 30min. Following blocking using blocking buffer (5% non-fat milk prepared in TBS-T [10mM Tris-HCl, pH 7.4, 150mM NaCl, 0.05% [v/v] Tween 20]) for 1h, the membrane was probed with an anti-EFA6R rabbit polyclonal antibody (1:500) (8) or an anti-β-actin mouse monoclonal antibody (1:10,000) (R&D Systems, UK) diluted in blocking buffer. After washing 3 times with TBS-T, the membrane was incubated with horseradish peroxidase (HPR)-conjugated secondary antibody (GE Healthcare, USA) diluted 1 in 2500 in blocking buffer. The membrane was incubated in the ECL Select substrate (GE Healthcare Life Sciences, UK) and visualised bands using a ChemicDoc XRS system (Bio-Rad, UK) as described (26). The ovarian OncoPair INSTA-Blot (Novus Biologicals, UK) was probed using the anti-EFA6R antibody or an anti-GAPDH goat polyclonal antibody (Everest Biotech, UK) diluted 1 in 1000 in blocking buffer.

### Immunohistochemistry

Immunohistochemistry was performed on commercially-obtained ovarian normal and cancer TMAs (US Biomax, USA) as described (24) and using a benchmark ultra IGC staining module (Roche/Ventana Medical Systems, USA). Briefly, following heat-induced antigen retrieval for 32min in CC1 retrieval buffer (pH 8.0 - 8.5), the anti-EFA6R antibody was used at a dilution of 1 in 150 and incubated at 36°C for 36min. OptiView HQ universal linker and HRP multimer were added for 8min to enhance stain quality. Diaminobenzidine (DAB) was used as the chromogen, and samples were counterstained with haematoxylin for 12min. The TMAs were blindly scored by 3 individuals. The protein expression was analysed by using a scoring method based on the sum of proportion of epithelial cells that showed staining (0 = none, +1 = <10%, +2 = 10-25%, +3 = 25-50%, +4 = 59-75%, +5 = 75-100%) and the intensity of staining (0 = none, +1 = weak, +2 = moderate, +3 = strong).

### Drug treatment

SKOV-3 cells were treated with indicated concentrations of 5-Aza-CdR (Sigma, UK) and SAHA (LC laboratories, UK) or 0.1% DMSO (solvent control). Drugs were replaced after 2 days of incubation. Following 4 days of the treatment, the cells were subjected to various functional assays.

### Functional assays

#### Cell viability

The viability of SKOV-3 cells treated with different concentrations of 5-Aza-CdR and SAHA for 4 days was assessed using the Kit-8 colourimetric cell viability kit (Biomake, USA).

#### Cell migration assay

SKOV-3 cells transfected without or with 100nM siRNA and simultaneously treated with 10μM 5-Aza-Cdr for 4 days were seeded into 2-well cell culture inserts (Ibidi, UK). After cell attachment (6h), the culture insert was removed and fresh medium added. Images of gap closure (migration) were recorded at 0h and 12h using an Olympus IX71 microscope and XM10 camera (Olympus, USA). The area between two edges of the migratory cells was measured using ImageJ software where cell migration presented as percentage of gap closure using the equation: ([pre-migration]_area_ - [post-migration]_area_/[pre-migration]_area_) x 100% (27).

#### Transwell migration/invasion assay

SKOV-3 cells transfected without or with siRNA and treated with solvent DMSO or 10μM Aza-Cdr for 4 days, or cells transfected with various EFA6R constructs DNA for 3 days, were resuspended in RPMI medium and seeded (250μl) into a 0.8μm pore sized polycarbonate membrane ThinCert™ cell culture insert (Greiner Bio-one, UK), pre-coated with 100μl of 1.2mg/ml of Matrigel (Sigma, UK) (for cell invasion assay), placed in a well of a 24-well plate. RPMI medium containing 10% FBS (as the chemoattractant) was placed outside of the insert within the well. Following 8h of cell migration or 16h of cell invasion, the inserts were washed twice with PBS, fixed with 4% paraformaldehyde (PFA) (Sigma, UK) for 10min, and stained with 0.4% crystal violet (Sigma, UK) for a further 10min. The staining was extracted using 5% SDS and absorption was subsequently measured at 570 nm using a plate reader.

#### Immunofluorescence

This was carried out as previously described (8). SKOV-3 cells grown on glass coverslips were transfected with GFP-EFA6R constructs for 48h, fixed with 4% PFA for 15min and incubated with 1μg/ml DAPI (Sigma, UK) in PBS for 5min to stain the nucleus. Coverslips were mounted on glass microscope slides using mounting solution (0.1M Tris-HCl, pH 8.5, 10% Mowiol [Sigma, UK] and 50% glycerol) containing 2.5% DABCO (1,4 diazabicyclo(2.2.2)octane) (28) and cells imaged using LSM710 confocal fluorescence microscope (Carl Zeiss AG, Germany) with a 63x oil-immersion objective lens. Scale bar in confocal images represents 10μm. The confocal images shown are representative of >50 cells from at least three independent cell preparations.

### Data Analysis

Data were analysed using GraphPad Prism software (USA). In some selected immunoblotting images, the average of three independent experiments is displayed as fold-change or % change of expression. A value of *P* >0.05 was considered not significant (ns) whereas *P* <0.05, *P* <0.01, *P* <0.001 and *P* <0.0001 (denoted as *, **, *** and ****) were used as the general limit of significance.

## Acknowledgements

We are grateful to Prof. Deyarina Gonzalez (Swansea University, Swansea, UK) for providing T1074 and SKOV-3 cell lines, Dr Marion Curt (University of Chicago, Chicago, IL, USA) for providing OVSAHO and Caov-3 cell lines and Dr James Cronin (Swansea University, Swansea, UK) for providing OVCAR8, COV504 and COV318 cell lines. We also thank members of the VK’s lab for helping by providing various reagents necessary for the study.

## Author contributions

V.K. conceived the project, designed and performed experiments and reviewed the manuscript. S.T-J. performed and analysed experiments and wrote the manuscript. W.W: analysed the data and reviewed the manuscript. K.M performed experiments and reviewed the manuscript. All authors had final approval of the manuscript.

